# Subtle changes in crosslinking drive diverse anomalous transport characteristics in actin-microtubule networks

**DOI:** 10.1101/2020.12.01.405142

**Authors:** S. J. Anderson, J. Garamella, S. Adalbert, R. J. McGorty, R. M. Robertson-Anderson

**Author notes:** Equal contributions (R. J. McGorty, R. M. Robertson-Anderson).

## Abstract

Anomalous diffusion in crowded and complex environments is widely studied due to its importance in intracellular transport, fluid rheology and materials engineering. Specifically, diffusion through the cytoskeleton, a network comprised of semiflexible actin filaments and rigid microtubules that interact both sterically and via crosslinking, plays a principal role in viral infection, vesicle transport and targeted drug delivery. Here, we elucidate the impact of crosslinking on particle diffusion in composites of actin and microtubules with actin-actin, microtubule-microtubule and actin-microtubule crosslinking. We analyze a suite of complementary transport metrics by coupling single-particle tracking and differential dynamic microscopy. Using these orthogonal techniques, we find that particles display non-Gaussian and non-ergodic subdiffusion that is markedly enhanced by cytoskeletal crosslinking of any type, which we attribute to suppressed microtubule mobility. However, the extent to which transport deviates from normal Brownian diffusion depends strongly on the crosslinking motif – with actin-microtubule crosslinking inducing the most pronounced anomalous characteristics – due to increased actin fluctuation heterogeneity. Our results reveal that subtle changes to actin-microtubule interactions can have dramatic impacts on diffusion in the cytoskeleton, and suggest that less mobile and more locally heterogeneous networks lead to more strongly anomalous transport.

## Introduction

The cytoskeleton is a complex network of filamentous proteins, including semiflexible actin filaments and rigid microtubules^1–3^. Numerous crosslinking proteins that can link actin to actin^4–7^, microtubules to microtubules^8–10^, and actin to microtubules^11–15^ enable the cytoskeleton to adopt diverse architectures and stiffnesses to drive key processes such as cell motility, meiosis and apoptosis^10, 16–18^. These varying structural and rheological properties, in turn, directly impact the intracellular transport of vesicles and macromolecules traversing the cytoplasm^14,15,19,20^. More generally, thermal transport of particles in biomimetic, cell-like and crowded environments continues to be intensely investigated due to the intriguing anomalous properties, i.e. properties not found in normal Brownian motion^14,15,19–24^, that have been reported in these systems.

Single-particle tracking (SPT), often used to characterize particle diffusion in complex environments, can determine anomalous transport characteristics such as: subdiffusion, in which the mean-squared displacement (MSD) scales as ~Δ*t*^*α*^ where *a* < 1^23, 25–27^; ergodicity-breaking, in which the time-averaged MSD differs from the ensemble-averaged MSD^28–30^; and non-Gaussianity, in which the distribution of particle displacements deviates from the normal distribution consistent with Brownian motion^31–36^. While single particle tracking measures the motion of individual particles to measure transport properties, differential dynamic microscopy (DDM) extracts complementary information inaccessible to SPT by probing the ensemble scale. Anomalous transport dynamics manifest in DDM measurements in the form of intermediate scattering functions (ISF) fit by stretched exponentials^37–39^ or ISFs that do not decay to zero, but have a non-zero plateau^39–41^ – hallmarks of heterogeneity and nonergodicity, respectively.

We previously showed that varying the relative concentrations of actin and microtubules in entangled actin-microtubule composites (without crosslinkers) led to surprising and complex effects on particle transport in these networks^15^. Namely, as the ratio of actin to microtubules increased, transport became more subdiffusive, less ergodic and exhibited more pronounced non-Gaussianity. We attribute these results to the smaller mesh size of actin-rich networks compared to microtubule-rich networks. However, the ~100-fold higher rigidity of microtubules compared to actin filaments also affected the dynamics, leading to non-monotonic dependences of multiple transport metrics with respect to the ratio of actin to microtubules. These intriguing results beg the question as to the role that filament crosslinking, which directly impacts network rigidity and connectivity, plays on the transport of particles through cytoskeleton composites.

Here, we couple single-particle tracking (SPT) with differential dynamic microscopy (DDM) to elucidate the anomalous transport of microspheres through crosslinked cytoskeleton composites over a broad spatiotemporal range. To systematically determine the role of crosslinking on particle transport, we examine co-entangled actin-microtubule composites in which we fix the concentration of actin, microtubules and crosslinkers and vary the type of crosslinking to include actin crosslinked to actin (A-A), microtubules crosslinked to microtubules (M-M), actin crosslinked to microtubules (A-M), both actin crosslinked to actin and microtubules crosslinked to microtubules (A-A/M-M), and compare to networks without crosslinking (None) (Fig. 1). We find that transport in all networks is subdiffusive, non-Gaussian and non-ergodic – as measured by both SPT and DDM. Further, crosslinking appreciably enhances subdiffusion, ergodicity-breaking and non-Gaussianity compared to unlinked networks. However, the degree to which each parameter is enhanced depends on the crosslinking motif, with networks with actin-microtubule crosslinking (A-M) inducing the strongest anomalous transport features and networks with actin-actin and microtubule-microtubule crosslinking (A-A/M-M) inducing the weakest. These findings dovetail with our previous work, wherein we show that the type of crosslinking causes surprising and distinct changes to actin and microtubule mobility^14, 42–44^. Taken collectively, our results indicate that the effect of crosslinking on transport is dictated by the suppression of microtubule mobility, whereas the relative impact of the crosslinking type on transport is modulated by the heterogeneity and rate of actin mobility.

**Figure 1.**
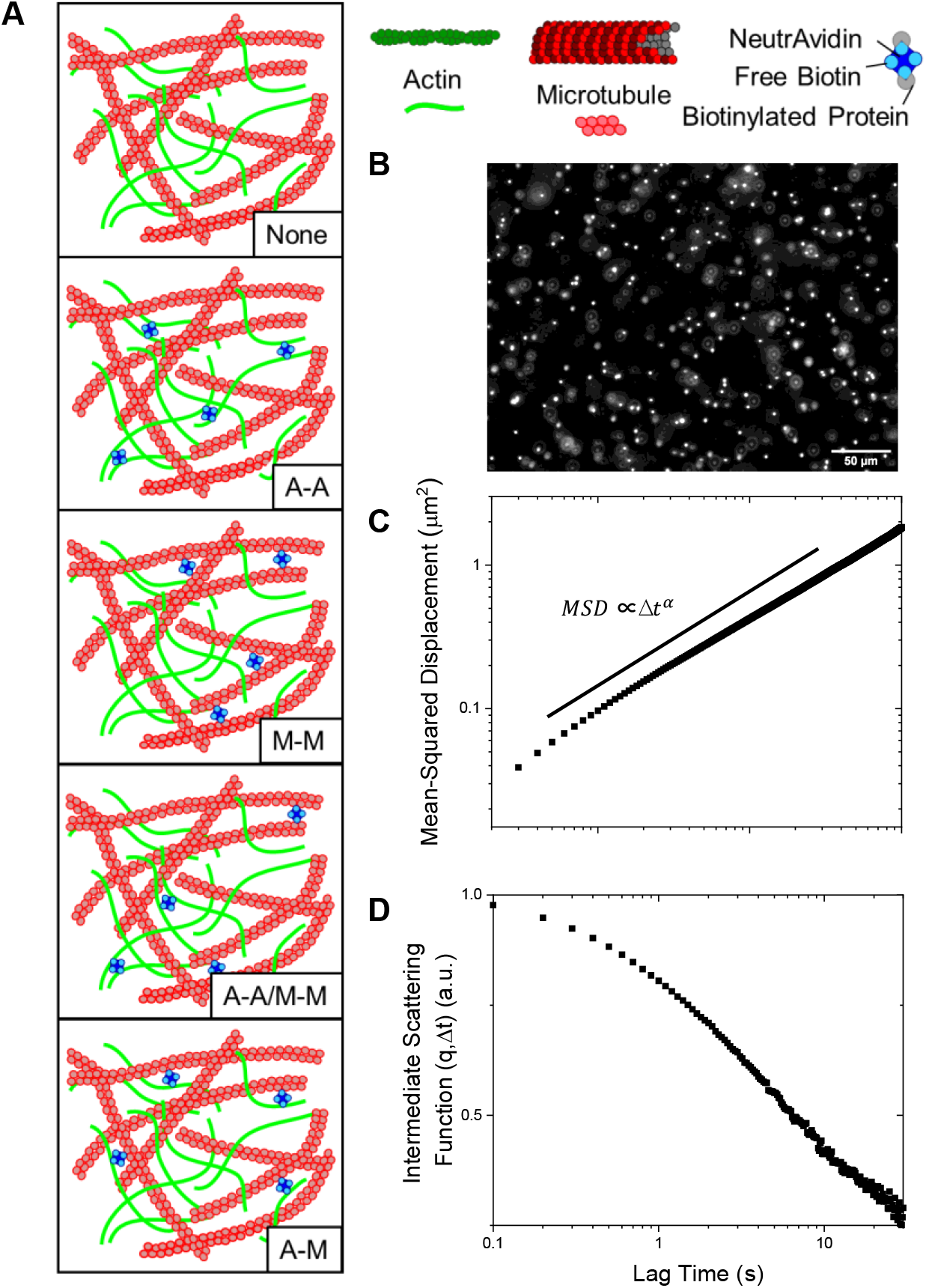
Experimental approach to examine the impact of crosslinking on anomalous transport in cytoskeleton networks. (A) Schematic of the different crosslinking motifs created in actin-microtubule networks: no crosslinkers (None), actin crosslinked to actin (A-A), microtubules crosslinked to microtubules (M-M), both actin-actin and microtubule-microtubule crosslinking (A-A/M-M), and actin crosslinked to microtubules (A-M). Biotinylated actin filaments and/or microtubules are crosslinked with NeutrAvidin to achieve the different motifs. (B) Videos of diffusing 1 μm fluorescent microspheres are collected and analyzed using single-particle tracking (SPT) (C) and differential dynamic microscopy (DDM) (D). (C) For SPT, the mean squared displacement (MSD) is plotted versus lag time (*Δt*) and fit to the power-law function MSD ∝ (Δ*t*)^*α*^ that describes anomalous diffusion. (D) For DDM, intermediate scattering functions are generated and fit to 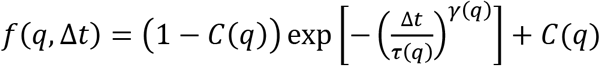 to extract the decay times *τ*(*q*), stretching exponents *γ*(*q*), and nonergodicity parameters *C*(*q*) for each condition. Data shown in (C) and (D) are for particles diffusing in the network without crosslinking (None).

## Results

We characterize the thermal transport of microscopic particles diffusing in cytoskeletal composites composed of co-entangled actin (A) and microtubules (M) with varying crosslinking motifs. We create composites with four different crosslinking motifs (A-A, M-M, A-M, A-A/M-M), as well as no crosslinking (None), utilizing single-particle tracking and differential dynamic microscopy to characterize the transport of microspheres diffusing in the various composites (Fig. 1).

In Figure 2A, we plot the mean-squared displacement as a function of lag time for particles diffusing in all five types of networks. As shown, all composites, with and without crosslinking, display anomalous subdiffusion, i.e. MSD ~ *Δt*^*α*^ where *α* < 1. However, all crosslinked networks are significantly more subdiffusive than the purely entangled case, especially at greater lag times. Further, in the crosslinked networks, the transport becomes more subdiffusive with increasing lag time, while in the unlinked network the degree of subdiffusion, as measured by *α*, is relatively constant. This effect is more readily shown by plotting the MSDs scaled by lag time (MSD/*Δt*) versus the lag time for each of the five composites (Fig. 2B). The solid lines in Fig. 2B act as a guide for *α* − 1 scaling, which is negative for subdiffusion.

**Figure 2.**
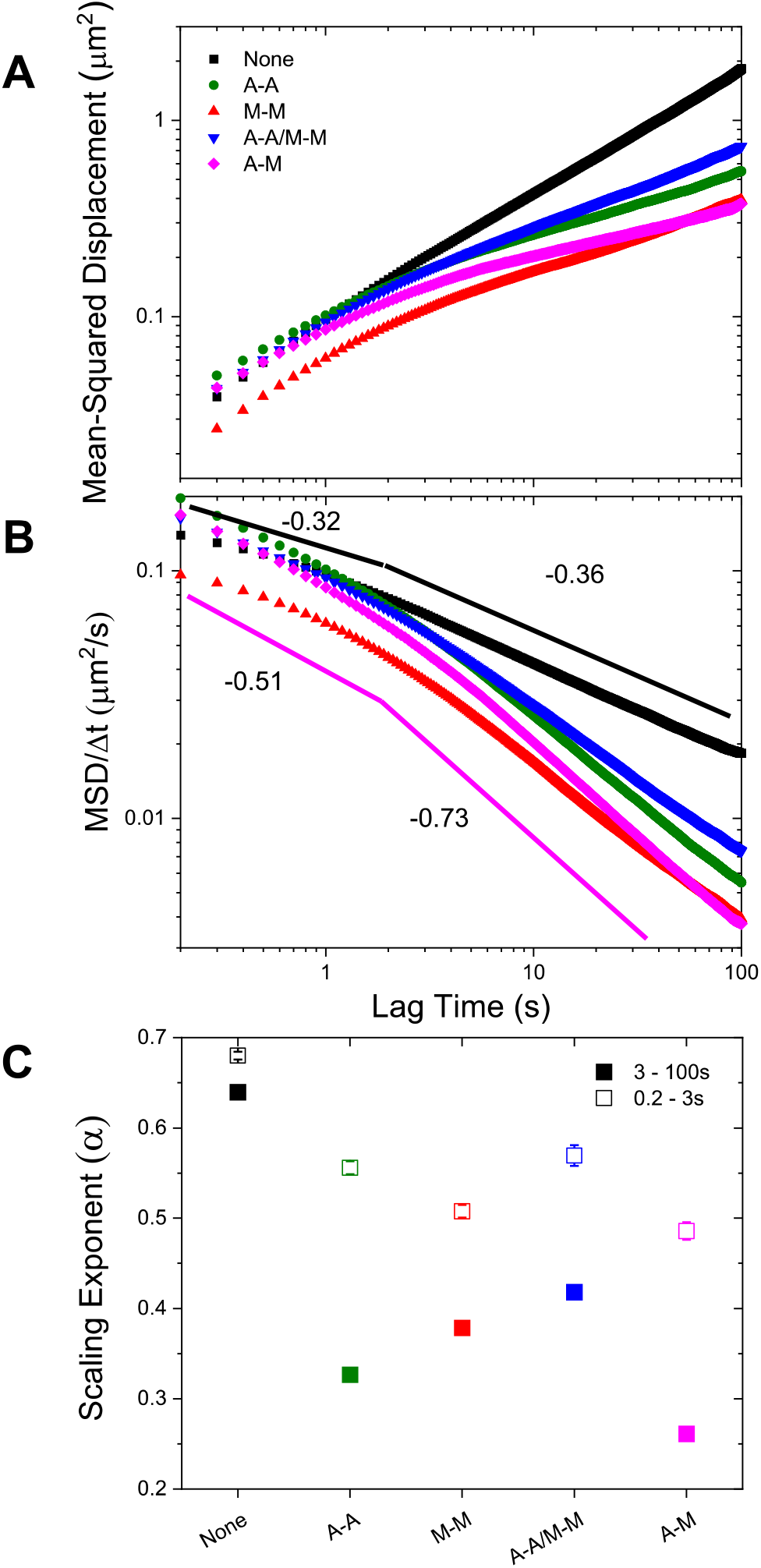
Crosslinking of cytoskeleton networks leads to multi-phase particle transport with more subdiffusive behavior at long times. (A) Mean-squared displacement (MSD) plotted as a function of lag time (Δ*t*) for each condition specified in the legend. (B) MSD scaled by lag time (MSD/*Δt*) versus lag time (*Δt*) for each condition. Pink and black lines are power-laws with scaling exponents (1– *α*) determined from fits to the A-M and None curves, respectively, for *Δt* = 0.2 – 3 s and *Δt* = 3 – 100 s. Steeper negative slopes indicate more subdiffusive transport. (C) Anomalous scaling exponents α from power-law fits of the MSDs. Open squares show the initial fitting region (0.2-3 s) and closed squares represent the long-time (3-100 s) fits. Error bars are the standard error calculated from four random subsets of data for each condition.

Due to the temporally variant characteristic of the subdiffusion for crosslinked networks, we fit the MSDs with power law functions (MSD ~ *Δt*^*α*^) over two distinct regions: 0.2–3 s and 3–100 s (Fig. 2C). For both time regimes, the non-linked composite exhibits a significantly higher scaling exponent than the crosslinked networks. Additionally, by quantifying the scaling exponent, it becomes clear that the non-homologous crosslinked network, in which actin and microtubules are crosslinked to each other (A-M), is the most subdiffusive over the entire spatiotemporal range. Conversely, the network in which both filaments are homologously crosslinked (A-A/M-M) appears to be the least subdiffusive, with scaling exponents that are higher than for networks in which only one filament type is crosslinked (A-A and M-M). Comparing the networks in which only one filament type is crosslinked, we find that microtubule-crosslinked networks (M-M) are more subdiffusive than actin-crosslinked networks (A-A), though this is less pronounced at long times. We note that for all networks, the molar ratio of crosslinkers to protein is held fixed. This is a critical detail, as it results in the actin and microtubules having twice as many crosslinkers per unit length in the A-A and M-M networks, respectively, as in the A-A/M-M network in which the crosslinkers are distributed among both filament types.

To examine potential mechanisms that give rise to the anomalous subdiffusion shown in Fig. 2, we evaluate the probability distributions of particle displacements, i.e. the van Hove distributions *G*(*Δx*, *Δt*), measured from SPT. Figure 3 shows van Hove distributions for a range of lag times from 0.3 s to 100 s for each network type. For reference, displacement distributions for particles undergoing normal Brownian motion are expected to be Gaussian^36^. However, as shown in Fig. 3, the distributions for all networks are distinctly non-Gaussian, with significant broad tails for large displacements. This phenomenon is commonly seen in crowded media in both synthetic and biological systems, and is a hallmark of dynamic heterogeneity^32–36,45,46^. However, in these systems, non-Gaussian transport is often transient and reverts to Gaussian at long enough times^47–49^. Conversely, we note the absence of a crossover to Gaussian transport in our systems at long lag times (100 s), suggesting that the dynamical processes governing transport in these networks have relaxation times longer than our measurement time scale. Although one may expect a reversion to Gaussian dynamics at long enough times, the time scales probed here are longer than the longest measured relaxation times of these cytoskeleton networks^6,43,50^.

**Figure 3.**
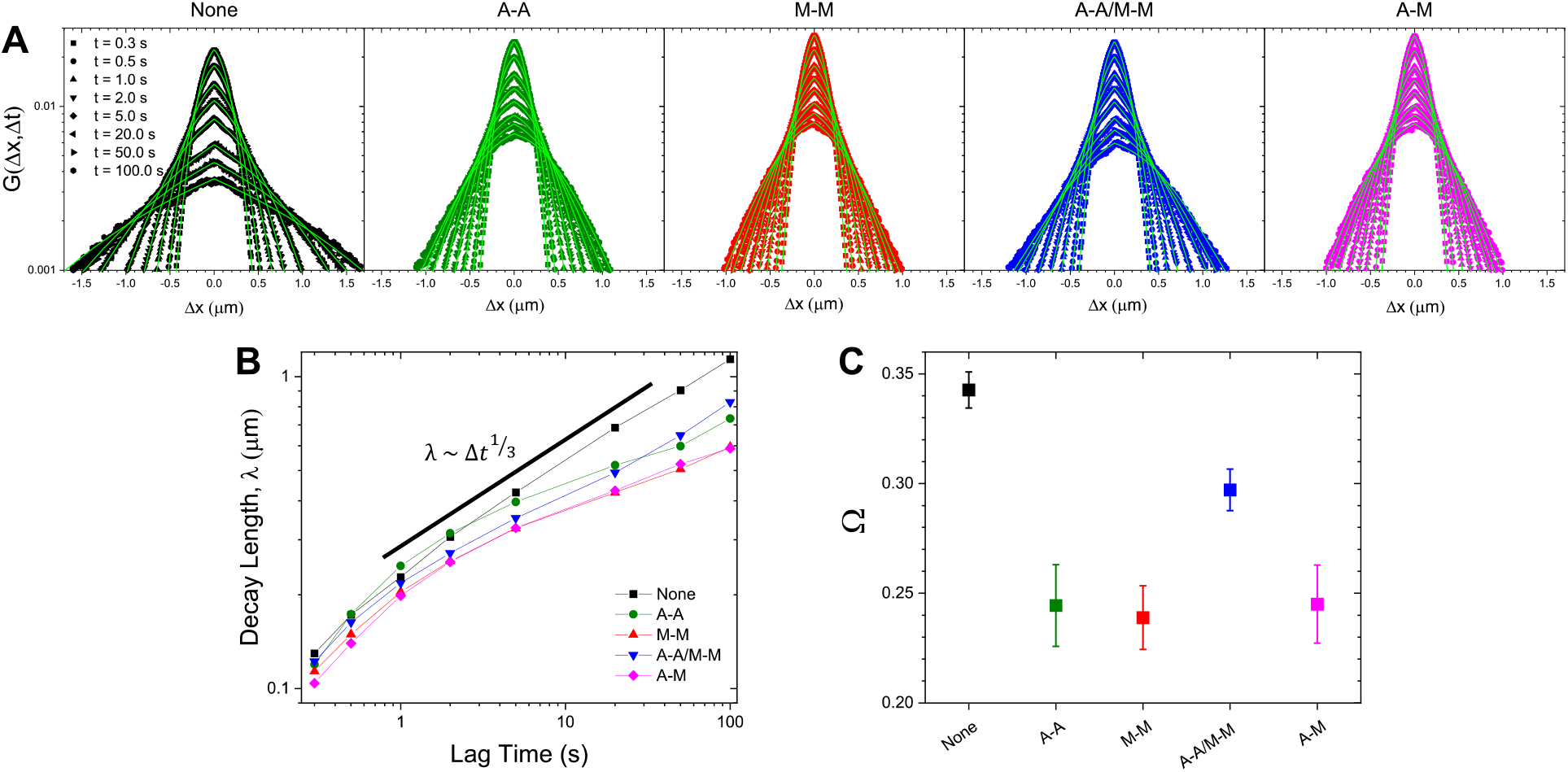
van Hove distributions reveal non-Gaussian, heterogeneous particle transport in all networks. (A) van Hove distributions (*G*(Δ*x*, Δ*t*)) for microspheres in composite networks without crosslinkers (None; black), with actin-actin crosslinking (A-A; green), microtubule-microtubule crosslinking (M-M; red), actin-actin and microtubule-microtubule crosslinking (A-A/M-M; blue), and actin-microtubule crosslinking (A-M; magenta) on a semi-log scale. Shown are the distributions for 0.3, 0.5, 1, 2, 5, 20, 50, and 100 seconds. The displacement distributions are distinctly non-Gaussian and were fit to a sum of a Gaussian and exponential distribution, 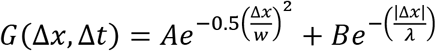, shown in bright green. (B) The characteristic length, *λ*, obtained from the exponential fits shown in (A), plotted against lag time. Points are connected with a line to guide the eye. (C) Scaling exponents, Ω, obtained from fitting the curves in (B) to a power-law *λ*(Δ*t*)~(Δ*t*)^Ω^.

We and others have previously found that non-Gaussian distributions in similar crowded and confined networks could be well-described by a sum of a Gaussian and exponential where the exponential, 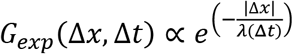, describes the large displacement tails^14,21,51^. The decay length *λ*(Δ*t*), is therefore best understood as the mean of length scales associated with the various relaxation processes that contribute to the exponential tails in the distributions. Moreover, this characteristic length has been shown to exhibit a power-law dependence on lag time with exponents of ~1/4 – 1/3^14,52^. To ascertain whether or not this decay length is sensitive to crosslinking motif, we plot *λ*(Δ*t*) against lag time and fit these curves to a power law, *λ*~Δ*t*^Ω^ (Fig. 3B). This decay length scales as a power law for all networks, though in the crosslinked networks the increase in the decay length with increasing lag time is smaller (smaller scaling exponent Ω). In order to quantify this scaling, we plot the scaling exponent Ω for each network type (Fig. 3C), which highlights the decreased growth of the characteristic length for the crosslinked networks relative to the unlinked network. Further, the decay length in the network with both filaments crosslinked (A-A/M-M) grows faster than in the homologous (A-A, M-M) or non-homologous (A-M) crosslinked networks (Fig. 3C). Finally, we note that the relevant time scales when the entangled and crosslinked networks diverge is strikingly similar to the time scales where the MSDs deviate from a single power law and become increasingly subdiffusive (Fig. 2).

To further investigate the network-dependent anomalous transport, we complement our SPT measurements with differential dynamic microscopy (DDM) analysis that we perform on the same samples we use for SPT. Specifically, we analyze the decay time, *τ*, of density fluctuations of an ensemble of particles across a range of spatial frequencies, *q*. As described in Methods, we fit the intermediate scattering function *f*(*q*, Δ*t*) to a stretched exponential while also taking into consideration the nonergodicity of the sample in order to extract dynamical information about the network. The stretching exponents (γ) we extract indicate the extent to which diffusion is anomalous. As seen in Figure 4, we find *γ* < 1 (~0.5 to 0.7) for all conditions, indicative of diffusion in confined media^40^. Further, the dependence of *γ* on the type of crosslinking follows a similar trend to that found for the SPT anomalous scaling exponent. Namely, the introduction of crosslinkers causes a significant decrease in the stretching exponent and non-homologous crosslinking leads to the smallest stretching exponent. While the smaller anomalous scaling exponent found in SPT analysis (Fig. 2) indicates more subdiffusion, a smaller stretching exponent represents a wider distribution of decay times that is indicative of more heterogeneity in the environment^37,38,40^. This trend supports our displacement distribution data (Fig. 3), which indicates substantial heterogeneity in all networks that is most apparent for A-M networks and least apparent for the non-crosslinked networks.

**Figure 4.**
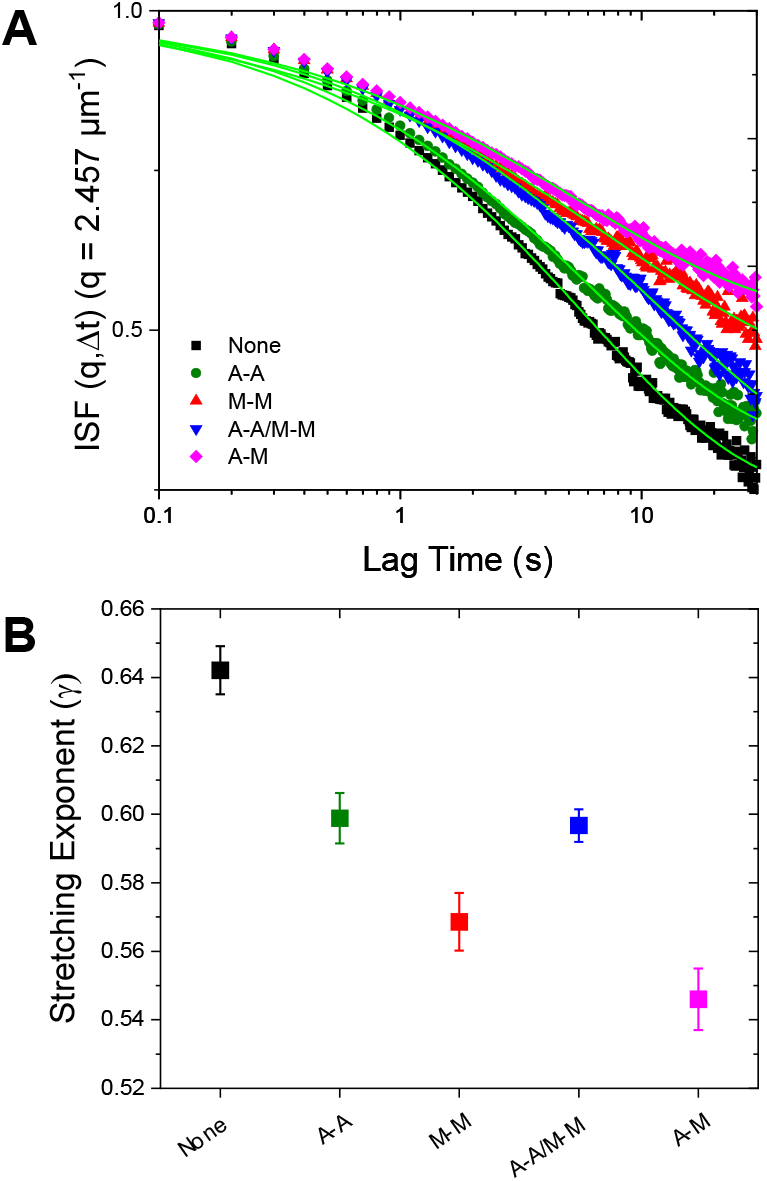
DDM analysis reveals heterogenous, non-ergodic transport amplified in crosslinked networks. (A) Intermediate scattering functions (ISF) are fit to a stretched exponential with a nonergodicity component, as described in Methods. The fits to the ISFs for each condition are plotted in bright green. The height of the long-time plateau reflects the nonergodicity of the transport. (B) DDM analysis shows increased local heterogeneity in crosslinked networks through decreasing stretching exponents (*γ*). The stretching exponent (*γ*) from the intermediate scattering functions for each condition is plotted. A stretching exponent less than 1 indicates the presence of heterogeneous crowded media. Error bars are the standard error calculated from the different videos taken for each sample.

Finally, to further quantify the extent to which the transport we report is anomalous, we evaluate three dimensionless parameters: the non-Gaussianity parameter *β*_*NG*_ determined via SPT data, the ergodicity-breaking parameter *EB* generated via SPT, and the non-ergodicity parameter *C* computed via DDM analysis (Fig. 5). For a particle undergoing Brownian motion, an ergodic process obeying Gaussian statistics, *β*_*NG*_ = *EB* = *C* = 0 for long times. As shown in Fig. 5A, *β*_*NG*_(Δ*t*) for all conditions decreases from an initial absolute maximum ~*O*(10) to a nearly time-independent plateau of ~*O*(1). However, the long-time plateau is significantly higher for the crosslinked networks relative the unlinked network, further evidenced by the time-average of *β*_*NG*_(Δ*t*) shown in the inset. We also compare the ensemble-averaged MSDs with the time-averaged MSDs through the ergodicity breaking parameter *EB*, computed via SPT and defined in the Methods. We observe that *EB* is non-zero for all conditions, indicating that the ensemble-averaged and time-averaged MSDs are not equivalent (Fig. 5B). Further, the non-homologously crosslinked network (A-M) is the least ergodic by a factor of ~2. The nonergodicity parameter, *C*, found independently through DDM analysis, shows strikingly similar trends to *EB* (Fig. 5B). *C* is a measure of the long-time plateau of the intermediate scattering functions and as such can vary between 0 and 1, with 0 indicating ergodic motion. Again, we find that the non-homologously crosslinked network (A-M) deviates the most from an ergodic process while the unlinked network deviates the least. The dependence of this deviation on the crosslinking motif follows a similar trend as our previously described parameters (subdiffusive scaling exponent, characteristic decay length and scaling, and DDM stretching exponent), wherein the unlinked network is consistently the least anomalous while the A-M network is the most anomalous.

**Figure 5.**
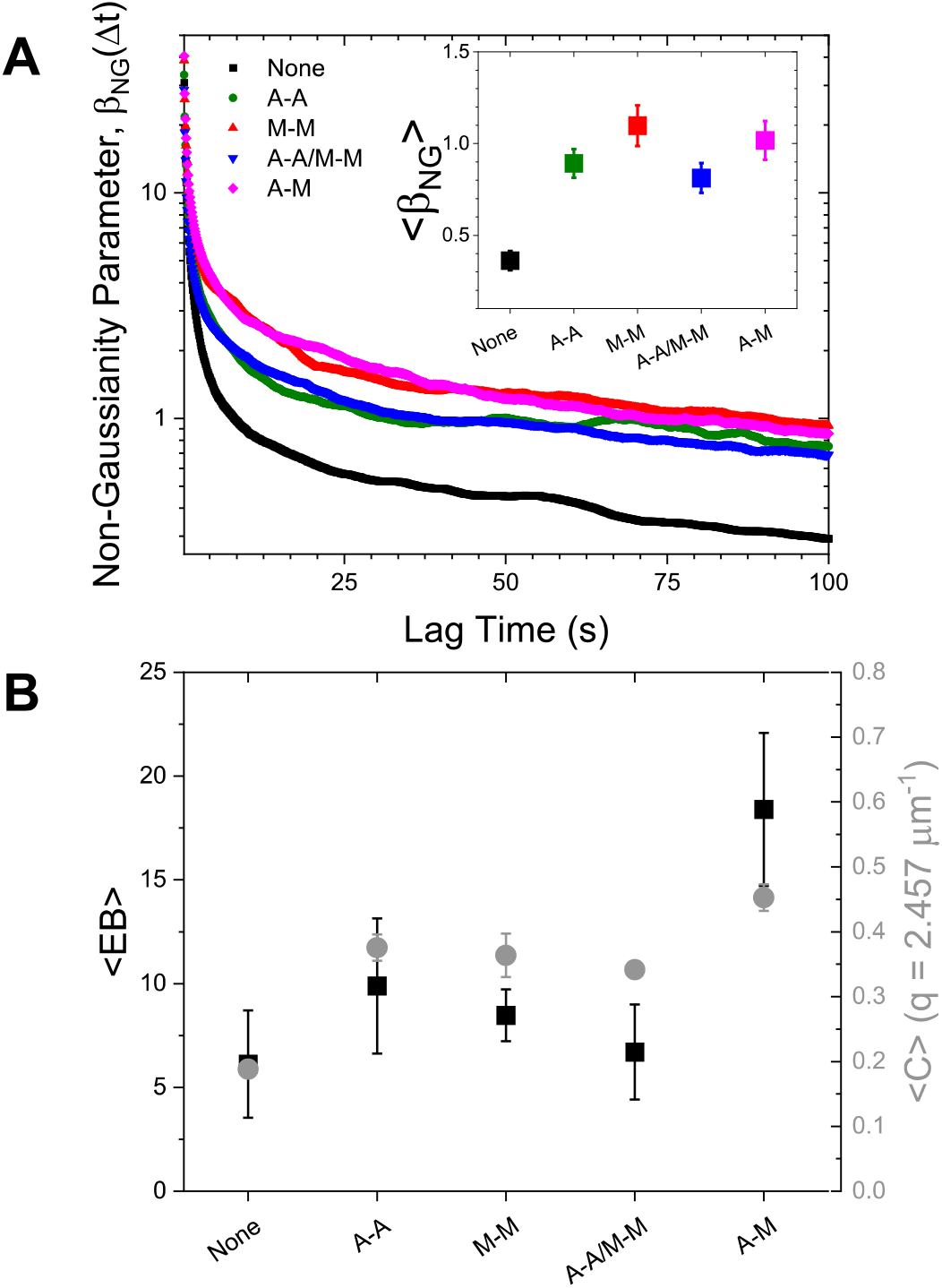
Metrics from both SPT and DDM show that crosslinking increases the non-Gaussianity and non-ergodicity of particle transport. (A) Non-Gaussianity parameter, *β*_*NG*_(Δ*t*), as function of time for each crosslinking type. Inset: Time-average of *β*_*NG*_. (B) Black, left axis: Time-average of the ergodicity breaking term *EB* as measured from SPT. Gray, right axis: Non-ergodicity parameter *C* as measured at wave vector *q* = 2.46 μm^−1^. Error bars represent the standard error calculated from the separate videos taken for each sample.

## Discussion

We use two independent measurements, SPT and DDM, to characterize the transport of microscopic particles in *in vitro* cytoskeleton networks with varying types of crosslinking. With these complementary techniques, we evaluate the degree to which the networks lead to deviations from normal Brownian diffusion with six different transport metrics to quantitatively characterize the network-dependent anomalous transport (Fig. 6). Collectively, our results indicate that particles undergo anomalous, non-Gaussian subdiffusion that is nonergodic in all networks. However, the degree to which these transport phenomena manifest is highly dependent on the type of crosslinking in the actin-microtubule composites.

**Figure 6.**
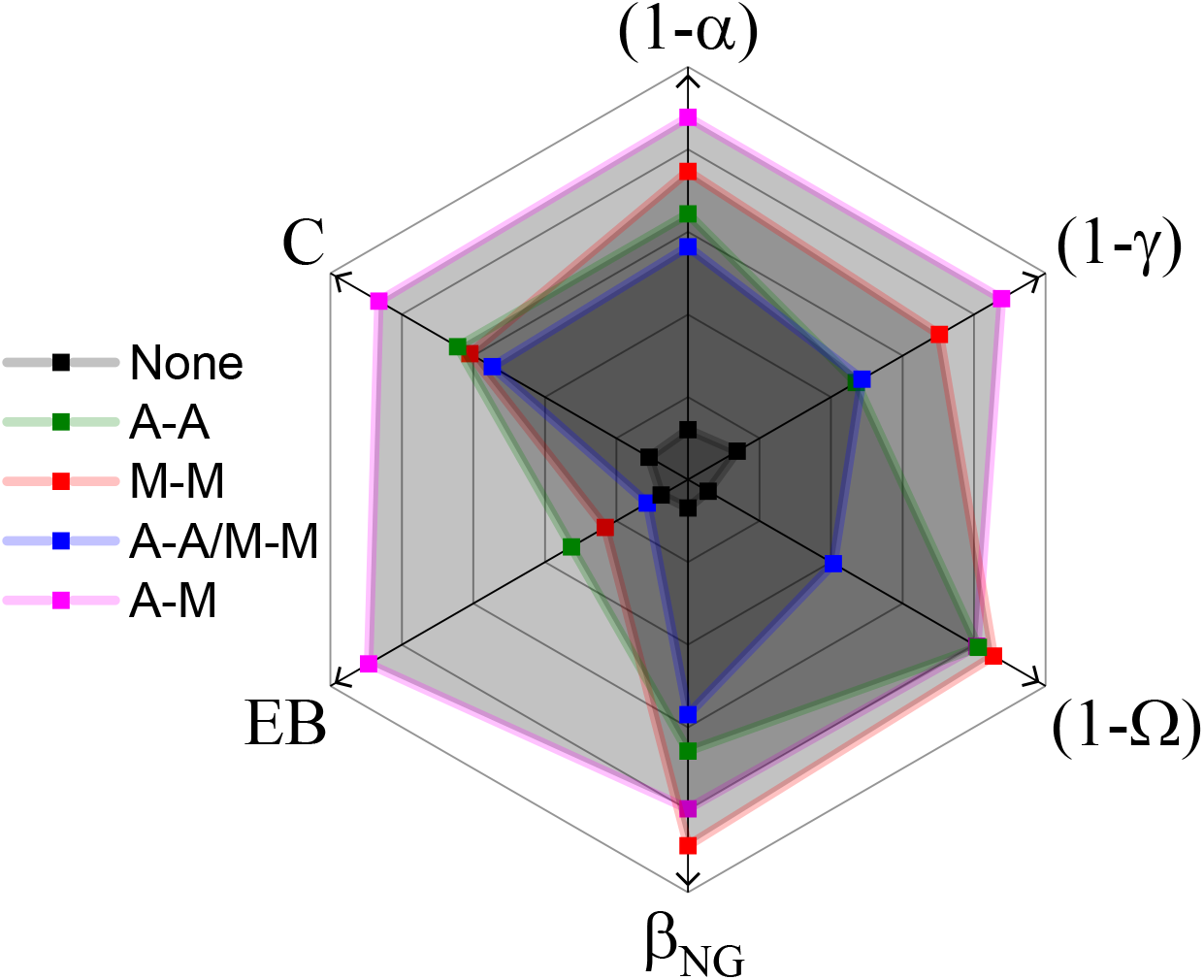
Multiple transport metrics highlight the degree to which crosslinking motif drives deviations from normal Brownian motion in composite cytoskeleton networks. Using the same color scheme as in previous figures, we show how the type of crosslinking, or lack thereof (black), influences subdiffusion (*α*), spatiotemporal heterogeneity (*γ*, Ω, *β*_*NG*_), and ergodicity (*EB*, *C*) across complementary, independent measurement techniques (SPT, DDM). A greater distance from the center (in the direction of the arrows) represents a greater deviation from normal Brownian diffusion. Each metric is scaled separately.

The unlinked network is consistently the least anomalous by all metrics (Fig. 6). While it may be intuitively expected that the least restricted network would have the least deviation from normal transport, this result is not trivial considering all networks have the same mesh size (ξ = 0.81 μm) due to the fixed molar concentrations of actin and microtubules. Further, the transport in the unlinked network is still complex, i.e. it is subdiffusive (Fig. 2), heterogeneous and non-Gaussian (Figs. 3, 4, 5), and exhibits ergodicity-breaking (Fig. 5). To understand the distinct transport characteristics in crosslinked versus unlinked networks we turn to our previous analysis of the filament dynamics in these networks^43^ which revealed that, upon crosslinking, the mobility of microtubules is significantly and equally suppressed by all crosslinking motifs, and the heterogeneity of microtubule fluctuations is likewise suppressed. This decrease in microtubule mobility likely acts to restrict or ‘cage’ the motion of the microsphere tracers to the local network mesh. This phenomenon is evidenced by the long-time limits in the MSD (Fig. 2A) and characteristic decay length *λ* curves (Fig. 3B). In nearly all crosslinked networks the particle MSDs do not exceed the square of the mesh size ξ^2^ and the decay lengths do not exceed the mesh size, indicating that trapping in the network mesh dominates transport in crosslinked networks, further evidenced by the decrease in proxies for ergodicity, *EB* and *C* (Figs. 5B, 6). Conversely, while transport in the unlinked network is indeed subdiffusive and non-Gaussian, MSDs exceed ξ^2^ and characteristic decay lengths, *λ*, are greater than ξ (Figs. 2A, 3B). This result suggests that the enhanced microtubule mobility in unlinked networks allows for faster network rearrangement to reduce caging, and instead couples microsphere transport to slow rearrangements of the local network^53^. This interpretation is bolstered by weaker ergodicity-breaking in the unlinked network compared to crosslinked networks (Fig. 5), as motion in a highly crowded, slowly evolving, biological environment without caging is an ergodic process^54^.

While we attribute the difference between crosslinked and unlinked networks to the suppression of microtubule fluctuations, this effect cannot explain the stark differences we find for transport among networks with varying crosslinking motifs. As shown in Figures 2–6, anomalous transport features are strongest for the non-homologous actin-microtubule crosslinked networks (A-M), followed by microtubule-microtubule (M-M), actin-actin (A-A) and A-A/M-M crosslinked networks. To understand this trend we look to our previous image analysis studies described above in which we found that, while microtubule mobility was equally reduced by all crosslinking types, the heterogeneity and rate of actin fluctuations were strongly dependent on the crosslinking type and displayed a similar trend to our transport metrics^43^. In particular, actin filaments displayed the slowest and most heterogeneous thermal fluctuations in A-M networks, followed by M-M and A-A networks, and the fastest and least heterogeneous fluctuations in A-A/M-M networks^43^.

Relative to the other crosslinked networks, transport within the A-A/M-M network is more weakly subdiffusive at long times and the temporal evolution of the decay length is steeper, suggesting that more homogenous and faster actin filament fluctuations lead to weaker deviations from normal Brownian motion. However, the universal decrease in microtubule mobility adds rigidity to the A-A/M-M network, such that particle caging still plays a significant role, leading to rises in non-ergodicity (*C*, *EB*), heterogeneity (Ω, *β*_*NG*_, *γ*) and subdiffusion (*α*) relative to the unlinked network.

## Conclusion

The cytoskeleton is a widely studied, complex network comprised, in part, of semiflexible actin filaments and rigid microtubules, along with myriad crosslinking proteins that crosslink actin to actin (e.g. alpha-actinin, filamin, spectrin, etc), microtubules to microtubules (e.g. MAP65, XCTK2, PRC1, etc) and actin to microtubules (e.g. MAP2, APC, proflin, plectin, etc). The diversity of crosslinking patterns possible with these crosslinkers not only directly alters the structure and dynamics of the cytoskeleton network, but, in turn, modulates the diffusion of biomacromolecules and particles in the cell. However, due to the complexity of the cytoskeleton, isolating the impact of different crosslinking motifs on diffusion through the cytoskeleton has proven difficult in vivo.

Here, we couple single-particle tracking with differential dynamic microscopy to elucidate the transport of microspheres in composite actin-microtubule networks in which we fix the concentrations of actin, microtubules and crosslinkers and only vary the type of crosslinking. Not only do the orthogonal techniques of SPT and DDM allow us to buttress our results between methodologies, but, more importantly, they capture transport properties over a spatiotemporal scale from single particles to the ensemble. Using these techniques, we have analyzed a robust suite of anomalous transport metrics to characterize particle dynamics in actin-microtubule networks with actin-actin, microtubule-microtubule and actin-microtubule crosslinking. We find that transport in all networks, even the network without crosslinkers, is subdiffusive, non-Gaussian and nonergodic. By introducing crosslinking, the transport becomes significantly more anomalous, per our metrics, which we argue arises from the significant decrease in the mobility of the microtubules, which, in turn, acts to increase the propensity for particle caging. The *type* of crosslinking also plays an important role in particle transport with actin-microtubule crosslinking resulting in the most extreme anomalous characteristics while networks in which both actin and microtubules are crosslinked to themselves but not each other exhibiting the least anomalous transport. Our previous results suggest that the dependence in transport characteristics on the crosslinking motif is a second order effect of varying actin mobility in the different networks. Namely, as the actin mobility decreases and its heterogeneity increases, the transport becomes more anomalous and increasingly controlled by the trapping of the particles within the network, as evidenced by the non-ergodicity and non-Gaussianity metrics.

Our intriguing results provide direct insight into particle transport in the cytoskeleton – important for processes such as viral infection, gene therapy, and drug delivery. More generally, our results have direct implications towards understanding macromolecular transport in biological crowded environments, polymeric materials and synthetic hydrogels. Finally, our measurement approaches, comprehensive suite of metrics, and well-characterized and controlled actin-microtubule composites serve as much-needed platforms for screening the transport properties of a wide array of tracer particles and network architectures.

## Methods

### Sample Preparation

Rabbit skeletal actin monomers and porcine brain tubulin dimers are purchased from Cytoskeleton (AKL99, T240) and suspended in G-buffer [2.0 mM Tris (pH 8), 0.2 mM ATP, 0.5 mM DTT, 0.1 mM CaCl_2_] and PEM-100 [100 mM piperazine-N,N′-bis(ethanesulfonic acid) (pH 6.8), 2 mM MgCl2, 2 mM EGTA], respectively. Resuspended actin and tubulin solutions are flash-frozen and stored at −80°C at concentrations of 2 and 5 mg/mL, respectively. To form crosslinked networks, we mix actin monomers and tubulin dimers at a 1:1 molar ratio in an aqueous buffer composed of PEM-100, 2 mM ATP, 1 mM GTP, 5 μM Taxol, and 0.05% Tween to a final protein concentration of c = 5.8 μM, as described previously^14,43,44,55^. To crosslink filaments, biotin-NeutrAvidin complexes with a 2:2:1 ratio of biotinylated protein (actin and/or tubulin) to free biotin to NeutrAvidin are preassembled and added to the solution at a crosslinker to protein molar ratio of *R*_cp_ = 0.02. We control the type of linking by varying the type(s) of biotinylated proteins (actin, tubulin, or both) we include in the crosslinker complexes^43,44^. For SPT and DDM measurements, we add a trace amount of 1 μm carboxylated fluorescent YG microspheres (Polysciences) which we coat with BSA to prevent nonspecific binding. We pipette final solutions into a sample chamber consisting of a glass slide and coverslip separated by ∼100 μm with double-sided tape. Chambers are sealed with epoxy and incubated at 37°C for 60 min to polymerize cytoskeletal proteins and form crosslinked networks.

### Imaging

For both single-particle tracking and DDM experiments, we image the microspheres using an Olympus IX73 inverted fluorescence microscope with a 20× 0.4 NA objective and a Hamamatsu ORCA-Flash 2.8 CMOS camera (180 nm/pixel). For SPT, we collect 16 1920×1440 pixel videos of 2000 frames at 10 fps for each condition. For each of the videos, we observe >40 trackable particles, producing a total of >600 particles tracked across two samples. For DDM, we collect 6 512×512 pixel videos of 5000 frames at 10 fps. Videos are analyzed by examining regions of interest (ROI) of 256×256 pixels.

### Single Particle Tracking

We use custom-written particle-tracking scripts (Python) to track the particle trajectories and measure the frame-to-frame x- and y-displacements (Δ*x*, Δ*y*) of the beads. From the displacements, we compute mean-squared displacements <Δ*x*^2^> and <Δ*y*^2^>. The average of <Δ*x*^2^> and <Δ*y*^2^> (MSD) as a function of lag time Δ*t* is fit to a power-law function MSD ∝ Δ*t*^*α*^ where α is the subdiffusive scaling exponent. For a system exhibiting normal Brownian diffusion, *α* = 1, while *α* < 1 indicates anomalous subdiffusion. Error analysis is performed by analyzing random subsets of 4-5 videos and calculating the standard error in α values from all 4 subsets.

We also evaluate probability distributions of the measured displacements (Δ*x*, Δ*y*) for various lag times (Δ*t*) to generate van Hove distributions (Fig. 3). These distributions are fit to a combination of a Gaussian and exponential function 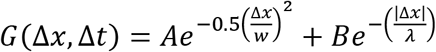, where *A* is the amplitude of the Gaussian, *w* is the Gaussian width, *B* is the amplitude of the exponential, and λ is the decay length. To further characterize the transport, we compute the non-Gaussianity parameter 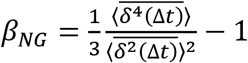 and the ergodicity breaking parameter 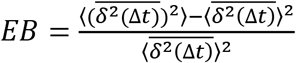 where δ^2^(Δ*t*) is the time-averaged MSD for the entire ensemble of trajectories^31,56^ (Fig. 5). For normal diffusion, both *β*_*NG*_ and *EB* trend towards zero, whereas anomalous transport manifests as *β*_*NG*_ > 0 and/or *EB* > 0.

### Differential Dynamic Microscopy

Following our previously described methods,^57,58^ we obtain the image structure function *D*(*q*, *Δt*), where *q* is the magnitude of the wave vector and Δ*t* is the lag time. The image structure function, or DDM matrix, can then be expressed as *D*(*q*, Δ*t*) = *A*(*q*)[1 − *f*(*q*, Δ*t*)] + *B*(*q*), where *A*(*q*) depends on the optical properties of the sample and microscope and *B*(*q*) is a function of the camera noise. The intermediate scattering function *f*(*q*, Δ*t*) is then described as

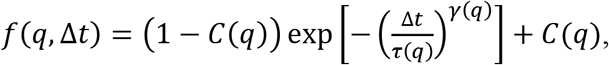

where *C*(*q*) is the nonergodicity parameter, *τ*(*q*) is the decay time, and *γ*(*q*) is the stretching exponent. Using methods similar to those described by Cho et al.^38^, we obtain both *A*(*q*) and *B*(*q*) prior to curve fitting. We find that *D*(*q*, Δ*t*) is independent of both *q* and Δ*t* in the highest *q*domains (*q* > 7.37 μm^-1^) as *A*(*q*) approaches zero. This is expected, as the value of *B*(*q*) is independent of *q* if camera noise is uncorrelated in space and time^59^. Thus, we take the minimum of the DDM matrix in this *q* regime and equate this value to *B*(*q*). To calculate *A*(*q*), we rely on the fact that, in linear space invariant imaging, the ensemble-averaged squared modulus of the Fourier-transformed images can be expressed as 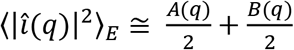, if the contributions from imperfections in the optical path are negligible to those in the sample^39^. Therefore, we calculate 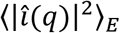 and, with *B*(*q*) already obtained, calculate *A*(*q*). In our previous work, we fit the image structure functions to extract the decay times and stretching exponents. Here, having obtained *A(q)* and *B(q)*, we fit the intermediate scattering functions directly to extract *C*(*q*), *τ*(*q*), and *γ*(*q*) using a combination of least squares and Levenberg-Marquardt curve fitting in Python.

## Data Availability Statement

The datasets generated and/or analyzed during the current study are available from the corresponding author upon request.

## Acknowledgements

This work was supported by NIH-NIGMS award no. R15GM123420 to R. M. R.-A. and R. J. M., as well as a William M. Keck Foundation Research Grant awarded to R. M. R.-A. R. J. M. acknowledges support from the Research Corporation for Science Advancement through the Cottrell Scholars program.

## Author Contributions

S. J. A. performed experiments, analyzed data, and wrote the manuscript. J. G. analyzed data and wrote the manuscript. S. A. analyzed data. R. M. R.-A. and R. J. M. conceived of and supervised the project and wrote the manuscript.

